# Prolonged Heat Treated Mesenchymal Precursor Cells Induce Positive Outcomes Following Transplantation in Cervical Spinal Cord Injury

**DOI:** 10.1101/2025.08.04.667962

**Authors:** Seok Voon White, Yee Hang Ethan Ma, Christine Plant, Alan R. Harvey, Giles W. Plant

## Abstract

Cellular transplantation therapies have been extensively used in experimental spinal cord injury research. However, there is no consensus as to what the most effective cellular controls for the therapeutic cell of interest are. For this reason, we examined if dead cells obtained through prolonged heat treatment can act as an appropriate cellular control for intravenously injected Sca-1+ mesenchymal precursor cells (MPCs) in C5 unilateral contusion cervical spinal cord injury. This was tested in single intravenous MPCs injection alone or intravenous MPCs plus intraspinal neural stem cells (NSCs) combinatory transplantation studies. MPCs were isolated from the compact bone of FVB mice while NSCs were isolated from the subventricular zone of luciferase-GFP transgenic FVB mice. Dead MPCs were obtained by heating at 72°C for at least 12 hours. In the MPCs only transplant study, injured mice received an injection of 1×106 dead or live MPCs D1 post-injury. Mice were then sacrificed at 8 weeks post-injury. In this study, intravenous injections of dead MPCs showed no statistical difference in injured paw usage compared to live MPCs, but behavior was improved compared to the media vehicle only control at D7 and D21. In the combinatory MPCs plus NSCs transplant study, injured mice received an intravenous injection of 1×106 dead or live MPCs D1 post-injury followed by intraspinal injection of 100,000 NSCs at D3 or D7 post-injury. Another two cohorts of mice received only NSCs at D3 or D7 post-injury. Mice were then sacrificed at 6 weeks post-injury. In this study, there was no functional difference in any of the groups in the dual injection study. Morphologically, mice receiving IV injection of dead MPCs had a smaller lesion size compared to the vehicular control, but the lesion size was larger than that of the lesion size in mice receiving live MPC injection. Dead cells elicited functional and anatomical benefits for the spinal cord injured mouse. In summary, dead cells obtained through prolonged heat treatment proved to be inconsistent and not optimal for use as cellular controls for cell transplantation studies in spinal cord injury but provides positive evidence for non-transplantation based cell therapies.

## 1. Introduction

Cellular transplantation remains at the forefront of experimental therapeutic strategies aimed at treating spinal cord injury (SCI). Current explored cell types include neural stem cells (NSCs) [1-3], neural precursor cells (NPCs) [4,5], hiPSC-derived mature neurons [6-9], regenerative peripheral cells such as Schwann cells (SCs) and olfactory ensheathing glia (OEGs) [10-12], and immune modulatory mesenchymal precursor cells (MPCs) [13-15]. These studies transplant live cells directly into the lesion via injections or through intra-venous injections. In some studies, dead cells are transplanted as a cellular control to the live cells, however, it is not known if dead cells by themselves would produce a therapeutic effect on the injured spinal cord. To better understand the mechanism and role of a transplanted cell type as a therapeutic intervention for SCI, it is important to understand the effects of both live and dead cells. Here, we have shown that dead mesenchymal precursor cells (dMPCs) can indeed show a therapeutic effect in the injured spinal cord and induce functional and anatomical improvements in a single IV injection and in combination with intraspinally injected NSC graft.

Previously, cell types of a different lineage were used as a cellular control in cell transplantation studies. For instant, olfactory ensheathing glia (OEG) and Schwann cells have been used as cellular controls for each other as they were deemed to be close in cell phenotype but functionally different. However, in transplantation for SCI, both OEG and Schwann cells have provided positive results when used as a treatment for SCI [16-21]. In addition, OEG were recently found to have the same neural crest origin as Schwann cells [22], which certainly complicates their use when interpreting the results if used as a cellular control for each other.

For MPCs, it had been suggested that dermal fibroblasts would be a good cellular control to examine their transplant efficacy. However, studies using dermal fibroblast transplantation have also yielded some positive outcomes, which questions the validity of these cells being an effective cellular control for MPCs [23, 24]. While certain cell types have been studied in great detail by the SCI research community and shown to elicit beneficial responses anatomically and/or functionally [25], even rarely studied cell types such as astrocytes [26], microglia [27] and macrophages [28] have been reported to elicit some positive outcomes. This indicates that a live cell of any kind is less than ideal as a cellular control for cell transplantation studies in SCI.

In our previous studies we used a vehicle, Hank’s buffered saline solution (HBSS), as a media only control, from here we refer to this as a vehicle control [1, 29]. A vehicle control only serves to ensure that the solution used for the injection does not by itself influence the injury. However, an injection of media only does not have the density, mechanical properties or substance of a cell suspension. Therefore, an additional control group to address the various factors is needed for cellular transplantation. Studies have shown that numerous techniques can be used to de-cellularize grafts such as peripheral nerves and skeletal muscle, including the use of chemical detergent [30-32], cold preserve protocol [33], freeze-thawing [34-36] and irradiation [37, 38].

Freezing tissue such as muscle [39] and peripheral nerve [40] is capable of killing all endogenous cells but leaves the basal lamina intact. These grafts are good controls for their experimental counterparts of muscle grafts and peripheral nerve loaded with Schwann cells, but identifying a control for the loaded cells is more difficult. Freeze-thawing fetal spinal cord tissue to produce a non-viable control graft for implantation has also been reported to be an adequate control for live fetal spinal cord tissue [41]. Heat treatment of muscle grafts has also been used to kill all endogenous cells, with the temperature used in the procedure shown to be crucial for a positive outcome [42].

In the present SCI study, we developed a protocol for obtaining dead MPCs (dMPCs) suspension and examined if the dead cells elicit a therapeutic effect to determine if they are valid cellular controls for transplantation of MPCs. In all the studies, unilateral C5 contusion SCI model was used. In the first study (referred to as IV study), IV injection of dMPCs was compared to IV injection of Hank’s Balanced Salt Solution (HBSS; vehicle control) and IV injection of live MPCs. All IV injections were done at D1 post-injury. In the second study (referred to as dual-transplantation study), we examined the integration and differentiation of intraspinal delivered NSCs at D3 or D7 post-injury when IV injection of MPCs or dMPCs was first delivered at D1 post-injury. Our overall aim was to determine if IV injection of dMPCs are suitable for cellular control for MPCs. The data from the studies showed that dMPCs elicited a positive effect behaviorally and anatomically when IV injected after SCI, albeit less effective than MPCs. Therefore, in the context of assessing MPC in spinal cord injury transplantation studies, it is important to note that dMPCs obtained through prolonged heating may not act as an appropriate inert cell control.

## 2. Materials and Methods

### Animals

Female FVB mice were used for all surgeries (12-14 weeks, Charles River). MPCs were isolated adult FVB mice. NSCs were isolated from adult transgenic mice that ubiquitously express green fluorescent protein (GFP) and firefly luciferase reporter genes (luc-GFP) driven by a chicken β-actin promoter [43, 44] with an FVB background (gift from Professor Joseph Wu, Stanford, USA) and FVB wild type mice. All animals were housed in a clean barrier facility on a 12/12-h dark-light cycle. The Stanford University Administration Panel on Laboratory Animal Care (APLAC) committee approved all protocols used in compliance with IACUC guidelines.

### Mesenchymal Progenitor Cell Cultures

MPCs were isolated from the compact bone as adapted from the method previously described [1, 29, 45]. Briefly, the ilium, femur and tibia were removed, cleaned and crushed to release the cells. The resulting cell suspension was depleted of CD5, CD45R, CD11b, Anti-Gr-1, 7-4, Ter-119 and CD3ε (Miltenyi Biotech, USA) positive cells using an autoMACs Pro Separator (Miltenyi Biotech, USA). The depleted cell population was then selected for Stem Cell Antigen-1 (Sca-1; Miltenyi Biotech, USA) in an autoMACs Pro Separator. Sca-1+ cells were plated down in media consisting of Minimum Essential Medium α (α-MEM) supplemented with 20% fetal calf serum, 1× GlutaMAXTM, 1× sodium pyruvate 100mM and 1 mg/ml gentamicin solution (MPC media; all reagents from Life Technologies, USA) at a density of 10,000 cells/cm2 and expanded until P4. MPCs were frozen at 1×106 cells/ml in MPC media with 7% dimethyl sulfoxide (DMSO) and stored in liquid nitrogen until required.

### Neural Stem Cell Culture

NSCs were isolated from the subventricular zone as adapted from Azari and colleagues [46]. Briefly, adult mice were euthanized with an overdose of sodium pentobarbital (Beuthanasia-D, 0.01ml/30g). Brains were removed and the thin layer of tissue surrounding the lateral wall of the ventricles was cut, being careful to exclude the striatal parenchyma and corpus callosum. The resulting tissue was minced and enzymatically digested with 3ml of 0.05% trypsin-EDTA (Life Technologies, USA) for 7 min in a 37°C water bath. The resulting cell suspension was mechanically dissociated using a pipette, and cells were washed and passed through a 40 μm cell strainer. Cells were seeded into one T25 flask per brain in 5ml of complete NSC medium supplemented with 20 ng/ml epidermal growth factor, 10 ng/ml basic fibroblast growth factor and 1μl/ml of 0.2% heparin (all items from StemCell Technologies, Canada) and expanded until P3 as free-floating neurospheres. Cells were then frozen per guidelines from StemCell Technologies. One flask was frozen per tube in 1.5 ml of media with 10% DMSO.

### Spinal cord injuries

Five mice were used per group for the IV study and ten mice were used per group for the dual-injection study. The injury model used in both studies was unilateral C5 contusion as previously described [1, 29]. Briefly, mice were anesthetized with isofluorane and a C5 laminectomy performed to expose the spinal cord. The Infinite Horizon Impactor (Precision Systems and Instrumentation, USA) with a custom-made 1 mm impactor head was used. An impact of 30 kDy with 3 sec dwell was used. Postoperative care consisted of subcutaneous administration of Pfizerpen (penicillin G potassium, 250,000 units/ml, Novaplus), buprenorphine (0.01 mg/kg, twice daily for 3 days) and saline (0.05 ml/g, twice daily for 3 days).

### MPCs Preparation for IV Injections

Previously frozen MPCs were rapidly thawed in a 37°C water bath and transferred to 15 mL tubes containing 5 mL of HBSS (Life Technologies, USA). Tubes were centrifuged for 5 mins at 400g and the resulting pellet resuspended with HBSS into a single-cell suspension. Excess DMSO was removed by repeating the previous step. The cell pellet was resuspended in 300μL of HBSS and transferred to a 1.5 mL tube and kept at room temperature before IV injection. Cell viability was checked using trypan blue with random samples taken before and after IV injections. Cells were >99% viable pre-injection and were >85% viable post-injection.

### dMPCs Preparation for IV Injections

A day before IV injection, MPCs previously frozen were rapidly thawed and prepared in a 1.5 mL microcentrifuge tube containing 300 μl of HBSS per 1×106 cells. The microcentrifuge tube was placed in a water bath at 72°C for over 12 hours (Figure 1A). The resulting cell suspension was tested for cell viability using trypan blue (Figure 1B). Viability of cells was confirmed to be <0.1% before being used for injections.

**Figure 1.**
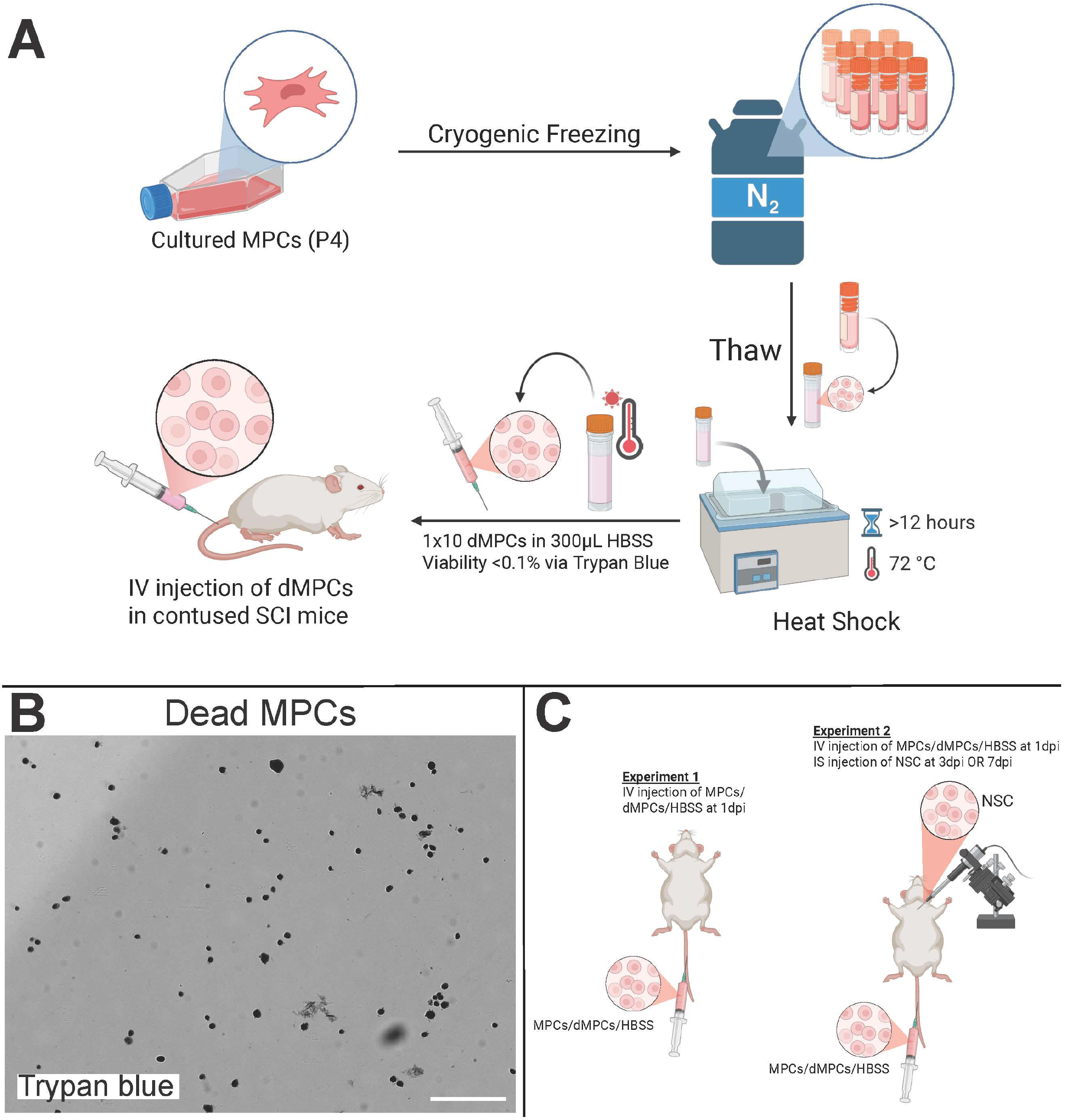
Experimental Schematics & Dead MPCs. (A) Schematic showing how dead MPCs were obtained for IV injection. (B) Dead MPCs as shown by trypan blue staining. Scale bar = 200 μm. (C) Illustration showing experiments 1 and 2. Briefly, in experiment 1, MPCs/dMPCs/HBSS were injected at 1 day post injury intravenously. In experiment 2, MPCs/dMPCs/HBSS were injected at 1 day post, and NSC were injected at 3 days or 7 days post injury intraspinally.

### Intravenous injections

MPCs or dMPCs were used for IV injections. Each mouse received 1×10^6^ MPCs or dMPCs in 300 μl HBSS per animal via the tail-vein. IV injection was carried out D1 post-injury. Vehicular control animals received 300 μl HBSS IV injection D1 post-injury. IV injection was carried out using a 1 ml syringe with 30 gauge needle.

### NSCs Preparation for intraspinal injections

Previously frozen NSCs were rapidly thawed in a 37°C water bath and transferred to 15 ml tubes containing 5 ml of HBSS. Tubes were centrifuged for 5 mins at 110g. The resulting cell pellet was resuspended in HBSS into a single cell suspension. Excess DMSO was removed by repeating the previous step. The resulting cell suspension was counted and resuspended in appropriate amounts of HBSS to ensure 100,000 cells/μl of HBSS. Trypan blue was used to check for cell viability before and after injections. Cells were >98% viable pre-injection, while post-injection the viability of the remaining cell suspension was >80%.

### Intraspinal Injections of NSCs

Mice receiving intraspinal injection of NSCs at D3 or D7 post-injury were anesthetized using isofluorane (2.5% in O2) and the spinal cord was exposed at the previous injury site. To prepare the spinal cord for injection, vertebra level C6 and C4 were clamped to straighten the spinal cord. A Nanoject IITM (Drummond Scientific, USA) with a custom glass pipette tip was used to inject 100,000 cells in 1 μl of HBSS at 200nl/min into 0.8 mm lesion epicenter. Postoperative care consisted of the administration of Pfizerpen (penicillin G potassium, 250,000units/ml), buprenorphine (0.01mg/kg, twice a day for 2 days) and saline (1ml/20g, twice a day for 3 days) subcutaneously.

### Group designation

After histological processing, mice that had injuries that crossed over midline were omitted from the overall results reported. The abbreviations used in text for each group as well as the total number of animals used per group for analysis (n) is shown in Table 1.

**Table 1.** List of experimental groups and abbreviations with numbers of animals per group.

| Group | Abbreviations | n= |
| --- | --- | --- |
| Intravenous HBSS D1 | HBSS | 5 |
| Intravenous MPC D1 | MPC | 5 |
| Intravenous dead MPC D1 | dMPC | 5 |
| Intraspinal NSC D3 | NSC_D3 | 8 |
| Intravenous MPC D1 with Intraspinal NSC D3 | MPC_D1/NSC_D3 | 9 |
| Intravenous dead MPC D1 with Intraspinal NSC D3 | dMPC_D1/NSC_D3 | 8 |
| Intraspinal NSC D7 | NSC_D7 | 8 |
| Intravenous MPC D1 with Intraspinal NSC D7 | MPC_D1/NSC_D7 | 9 |
| Intravenous dead MPC D1 with Intraspinal NSC D7 | dMPC_D1/NSC_D7 | 10 |

### Behavior

Mouse cylinder test for gross paw usage was used to collect behavioral data [47]. For the IV study, mice were recorded at D7, D21, D35 and D49 post-injury. For the dual-injection study, mice were recorded at D7, D14, D21 and D35 post-injury. Mice were placed in a Perspex cylinder and recorded for 5 mins. The number of left and right-paw touches was counted. Only full touches with hind leg rearing were counted. The percentage of right-paw usage was then calculated (right-paw usage/left + right-paw usage x 100%).

### Histology

For the IV study, mice were euthanized at D56 post-injury. For the dual-injection study, mice were euthanized at D42 post-injury. Mice were euthanized with an overdose of Beuthanasia-D and transcardially perfused with 4% paraformaldehyde. Spinal cord tissue was collected, post-fixed in 4% paraformaldehyde overnight and then 30% sucrose in PBS. Horizontal spinal cord sections were obtained using a freezing microtome (Leica, USA) at 50μm thickness. For the IV study, every one in four sections were mounted onto subbed slides and stained with Luxol Fast Blue with Cresyl violet counterstain. Neurolucida Neuron Tracing Software (MBF Bioscience, USA) was used to measure the lesion size. Lesion size was measured over five sections per mouse and then averaged as the total lesion area.

### Immunostaining

Prior to primary antibody incubation, tissue sections were blocked for at least 1 h at room temperature in diluent consisting of PBS + 10% normal donkey serum + 0.2% Triton X-100. Sections were incubated in primary antibody in diluent for secondary antibody in diluent for 30 minutes at room temperature. For the IV study, primary antibodies used were anti-βIII tubulin (1:800, Covance, USA), rabbit anti-glial fibrillary acidic protein (GFAP, 1:500, Dako, USA) and rabbit anti-laminin (1:800, Sigma-Aldrich, USA). Blood vessels were labeled with DyLight® 488 Lycopersion Esculentum (Tomato) Lectin (1:100; Vector Laboratories, USA). For the dual-injection study, primary antibodies used were βIII tubulin (mouse, 1:800; Covance), GFAP, anti-APC (mouse, 1:200; Abcam, USA) and green fluorescence protein (GFP, chicken, 1:10000; Aves Lab, USA). Sections were washed thrice in diluent before incubation in secondary antibody. The corresponding secondary antibody for each species was from Jackson Laboratories (USA) and included AF647 anti-mouse (1:800), DyLightTM 549 anti-rabbit (1:1000), and DyLightTM 488 anti-chicken (1:800). Confocal fluorescence images were obtained with a x20 objective on a Nikon C2 confocal using a 5×5 large image template scan with 3 μm z-stacks. Higher magnification images were taken with an x63 oil objective on a Nikon C2 confocal with 0.25 μm z-stacks.

### Dual-injection study: NSC counts

Transplanted GFP+ NSCs were counted in 8 sections per mouse. Using cellular morphology and antibodies labeling for βIII tubulin, GFAP and APC, the counted cells were categorized as neurons, astrocytes or oligodendrocytes

### Statistics

All one-way ANOVA with Tukey’s post-hoc test statistics were performed using GraphPad Prism Software. Values of P < 0.05 were considered statistically significant. All error bars are standard deviation.

## 3. Results

### IV study: Behavioral and lesion size outcome after IV injection of dead MPCs

The relative use of the injured paw (right) versus the uninjured paw (left) after cervical SCI was examined using mouse cylinder testing. There was no statistical significance seen between the right paw usages in the dMPC group versus MPC group at any of the measured time points (Figure 2A). However, there was statistical significance between the dMPC group compared to the HBSS at D7 (p=0.0308) and D21 (p=0.0263) (see asterisks, Figure 2A).

**Figure 2.**
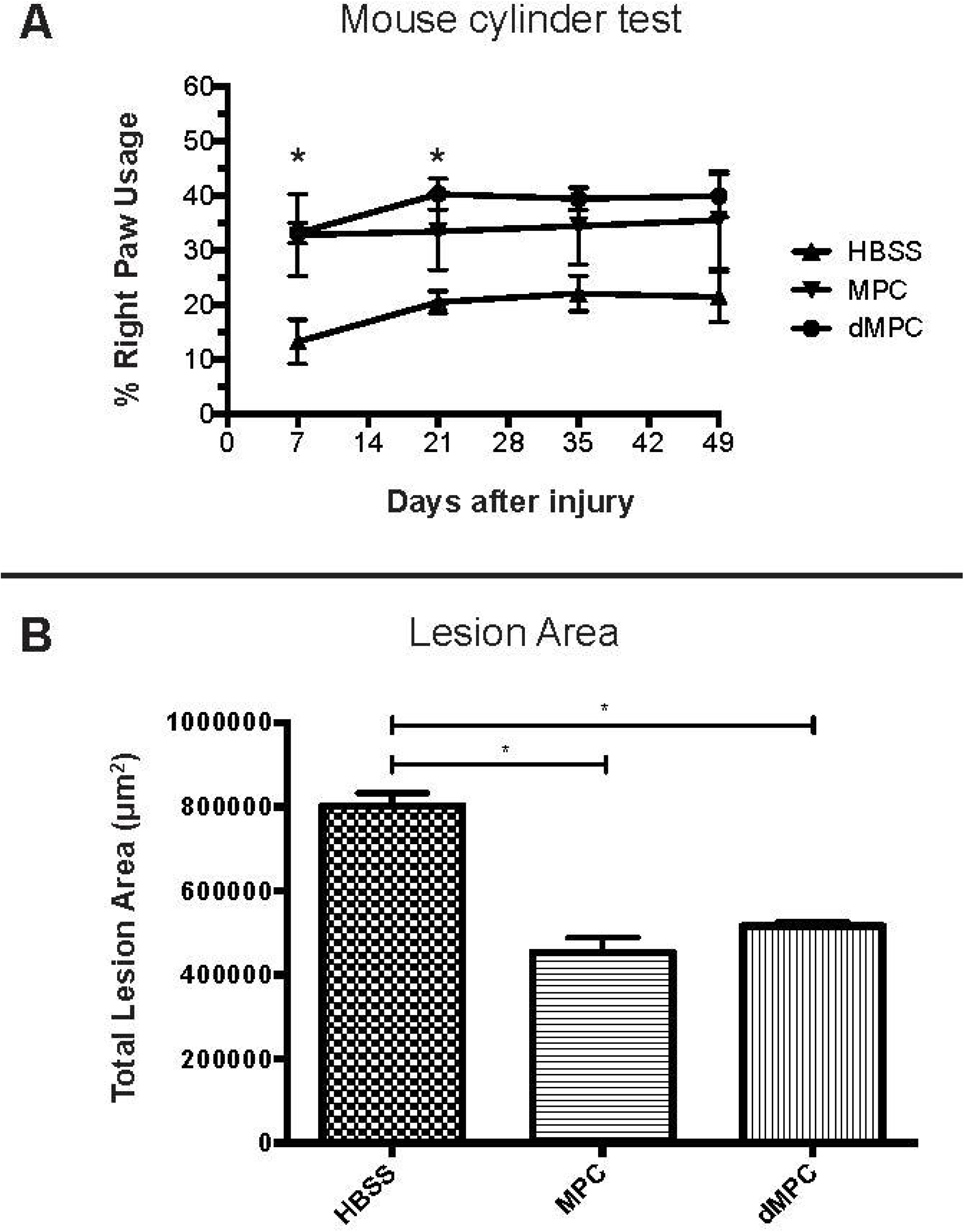
Behavioral and lesion size results for mice receiving IV injection of dMPC and MPC in cervical contusion SCI. (A) Mouse cylinder test results for mice with contusion injury receiving IV injection of HBSS, dead or live MPCs at D1 post-injury. There was no statistical difference between dead MPCs or live MPCs. The asterisk indicates statistical significance between the dMPC group and contusion + HBSS group at 7 and 21 days post-injury (p=0.0308 and p=0.0263 respectively). No statistical significance was seen at 35 and 49 days post-injury. (B) Lesion size results for mice with contusion injury receiving IV injection of HBSS, dead or live MPCs at D1 post-injury. There was no statistical difference in the lesion size at 8 weeks post-injury of mice receiving IV injection of dead MPC compared to mice receiving IV injection of live MPC D1 post-injury (p=0.6277). However, both groups showed statistically significant difference in the lesion size compared to the HBSS group (^*^p<0.0001).

Lesion size was measured using Neurolucida Neuron Tracing Software. There was no statistical significance seen in the lesion size when the dMPC group was compared to the MPC group (p=0.6277; Figure 2B). However, the lesion size in the dMPC group was, on average, approximately 100,000 μm2 larger than the lesion size in the MPC group (Figure 2B). There was statistical significance seen in the lesion size in the dMPC group compared to the HBSS group (p<0.0001). This was also seen in the MPC compared to the HBSS group (p<0.0001) (Figure 2B).

### IV study: Injury site profile after intravenous injection of dead MPCs

Spinal cord tissue sections were labeled with markers against βIII tubulin to examine axonal profile, GFAP to examine astrocytic profile, tomato lectin to examine vascularization profile and anti-laminin to examine extracellular matrix deposition profile (Figure 3 and 4). The axonal profile as determined by βIII-tubulin immunofluorescence in dMPC group showed some unorganized axonal sparing around the injury site compared to the MPC group (Figure 3Bii, Ciii). Within the injury site of the dMPC tissue, there were distinct areas that were devoid of any axons (see asterisks, Figure 3Bii). This was not observed in the MPC group but featured prominently in the HBSS group (see asterisks, Figure 3Aii, Cii). Using the astrocytic marker GFAP, we confirmed that the same area was also devoid of astrocytes. (Figure 3D, E). This was not observed in the MPC group (Figure 3F). Using anti-tomato lectin and anti-laminin, we observed massive blood vessel formation and extracellular matrix deposits at the lesion site in the HBSS group (Figure 4A, D). This was also seen to some extent in the dMPC group (Figure 4B, E) but not seen in the MPC group (Figure 4C, F) where the blood vessels and extracellular matrix deposits were less congregated and more organized. The histological observations indicate that the injury site in mice with dMPC injection was improved compared to mice with HBSS injection but not to the degree seen in mice with MPC injections.

**Figure 3.**
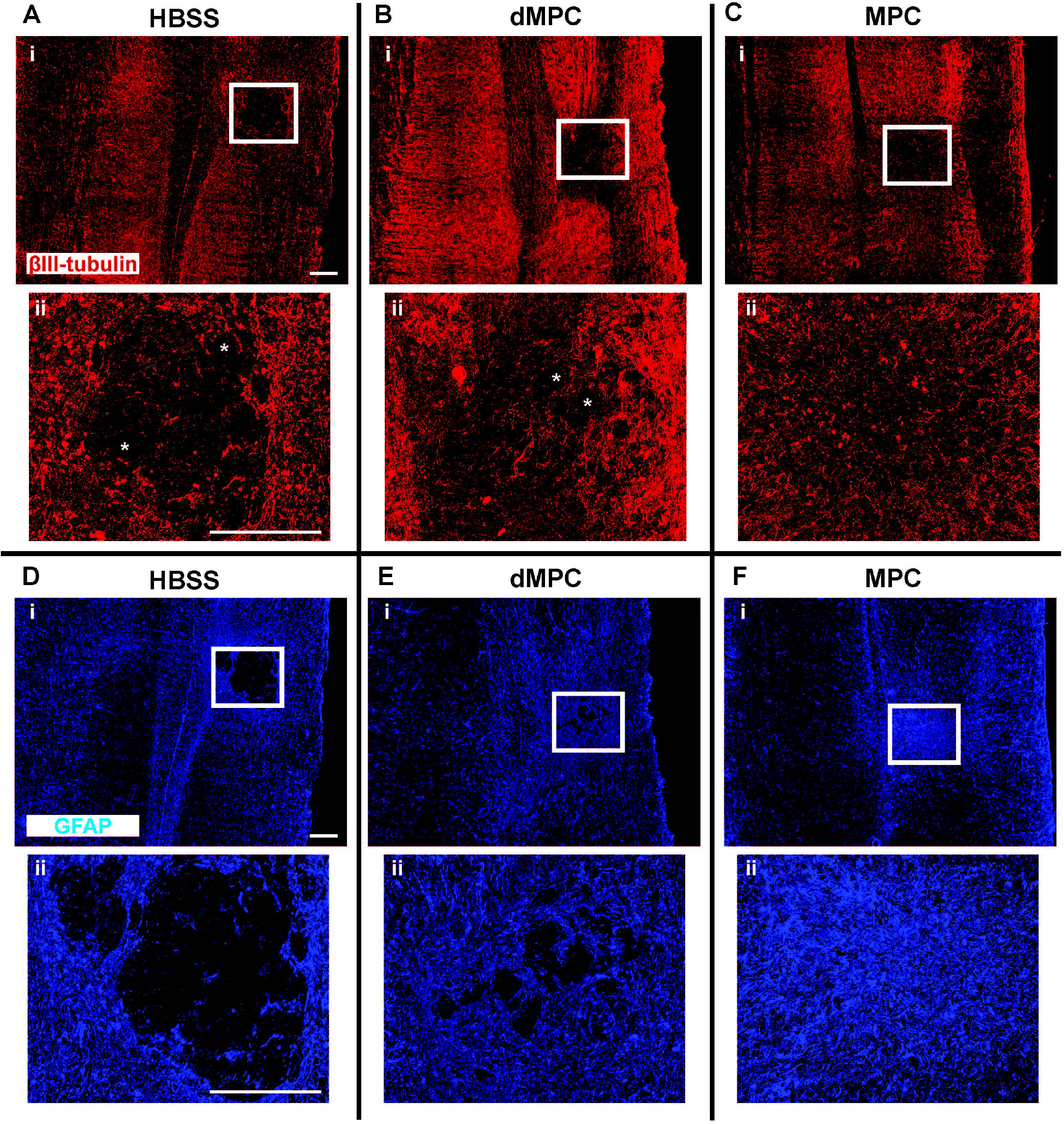
Axonal and glia profile 8 weeks post-injury. Images showing a stitched section of the spinal cord with the injury site labeled with βIII-tubulin+ axonal profile inmice receiving (Ai) HBSS, (Bi) dMPC and (Ci) MPC treatment. The corresponding (ii) images show the microscopic view of the injury site as outlined by the white boxes. Images showing a stitched section of the spinal cord with the injury site labeled with GFAP+ astrocytic glia profiles in mice receiving (Di) HBSS, (Dii) dMPC and (Cii) MPC treatment. The corresponding (ii) images show the microscopic view of the injury site as outlined by the white boxes. ^*^ shows areas devoid of axons. Scale bar = 200 μm.

**Figure 4.**
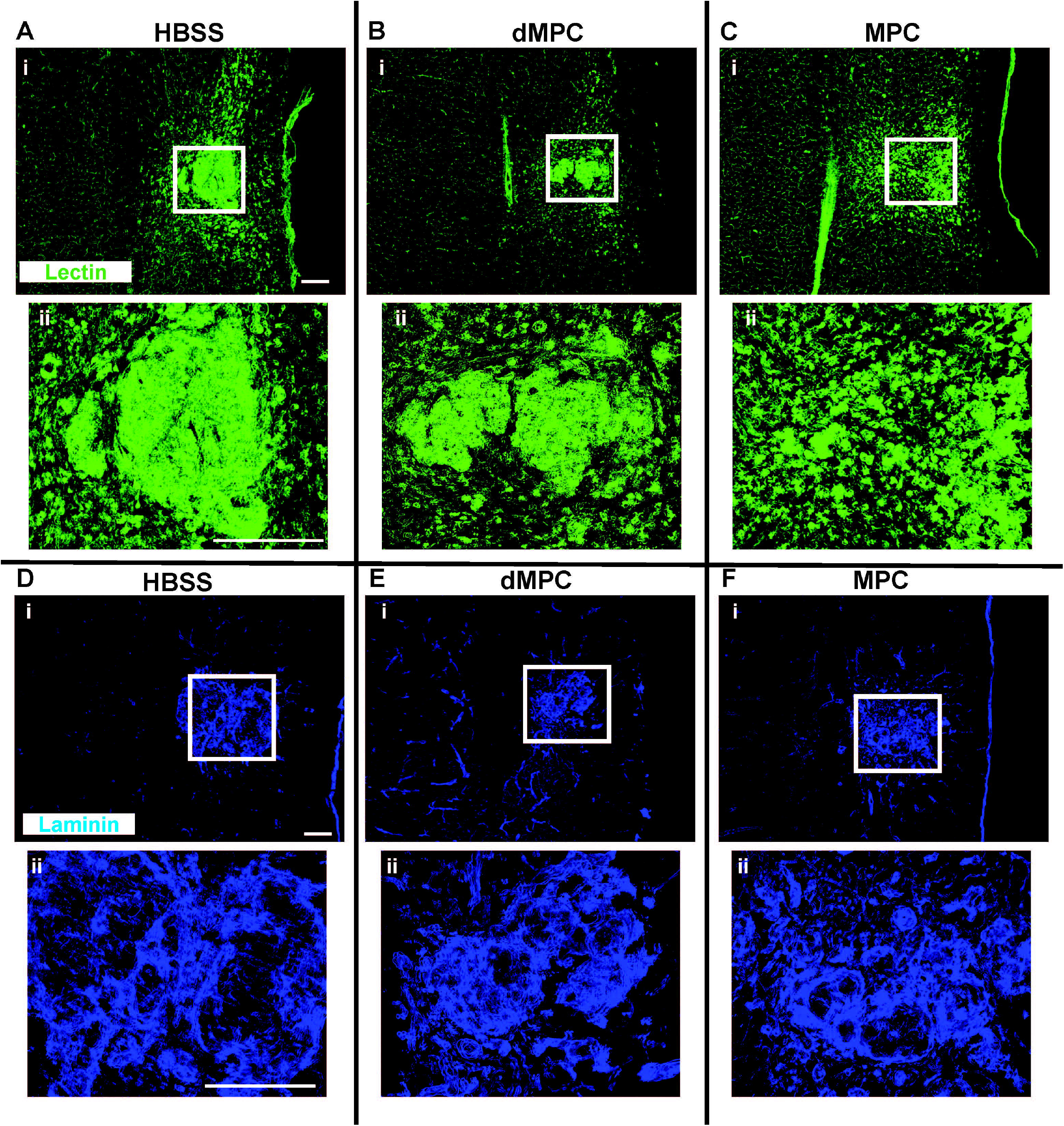
Vascularization profile 8 weeks post-injury. Images showing a stitched section of the spinal cord with the injury site labeled with lectin+ blood vessels in mice receiving (Ai) HBSS, (Bi) dMPC and (Ci) MPC treatment. The corresponding (ii) images show the microscopic view of the injury site as outlined by the white boxes. Images showing a stitched section of the spinal cord with the injury site labeled with laminin+ extracellular matrix profile in mice receiving (Di) HBSS, (Dii) dMPC and (Cii) MPC treatment. The corresponding (ii) images show the microscopic view of the injury site as outlined by the white boxes. Scale bar = 200 μm.

### Dual-transplantation study: Behavioral outcome after dual-injection of dMPCs with NSCs

Mouse cylinder behavioral test was used to examine right paw usage after IV injection of MPCs or dMPCs in conjunction with intraspinal NSC injection after cervical SCI. The results showed that there was no functional improvement in any of the treatments when compared to each other (Figure 5). This was regardless of whether NSCs were intraspinally injected at D3 (Figure 5A) or D7 (Figure 5B). Mice receiving MPCs injection prior to NSCs injection did not perform better compared to mice receiving dMPCs injection

**Figure 5.**
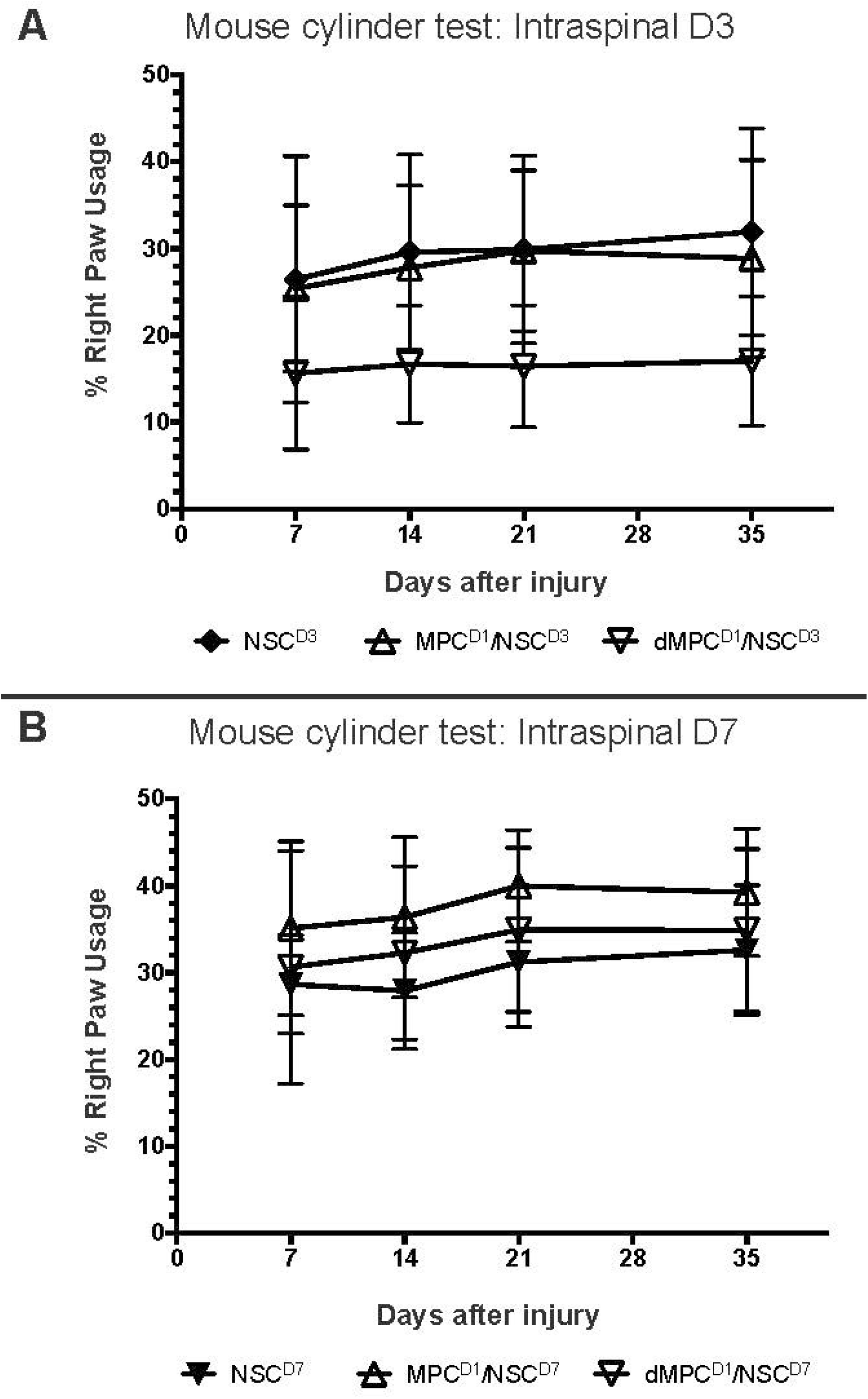
Behavioral results from mouse cylinder test for the dual transplantation study in mice receiving cervical contusion SCI. (A) Mouse cylinder test results for mice with cervical contusion injury receiving intraspinal injection of NSCs at D3 post-injury, with or without prior IV injection dead or live MPCs at D1 post-injury. There was no statistical difference between any of the groups at any of the time points examined. (B) Mouse cylinder test results for mice with cervical contusion injury receiving intraspinal injection of NSCs at D7 post-injury, with or without prior IV injection of dead or live MPCs at D1 post-injury. There was no statistical difference between any of the groups at any of the time points examined.

### Dual-transplantation study: Intraspinal injected NSCs survive and differentiate after intravenous injection of dead MPCs

Six weeks after the initial injury, mice were sacrificed and spinal cord tissue processed histologically with anti-GFP marker to identify for the transplanted NSCs. In all the groups receiving intraspinal injection of NSCs either D3 or D7 post-injury, a small amount of NSCs survived and differentiated into either astrocytes or oligodendrocytes (Figure 6). There was no evidence that any NSC differentiated into neurons. There was no statistical significance between the survival and oligodendrocyte differentiation of NSCs in the dMPC_D1/NSC_D3 or dMPC_D1/NSC_D7groups compared to the MPC_D1/NSC_D3 or MPC_D1/NSC_D7groups.

**Figure 6.**
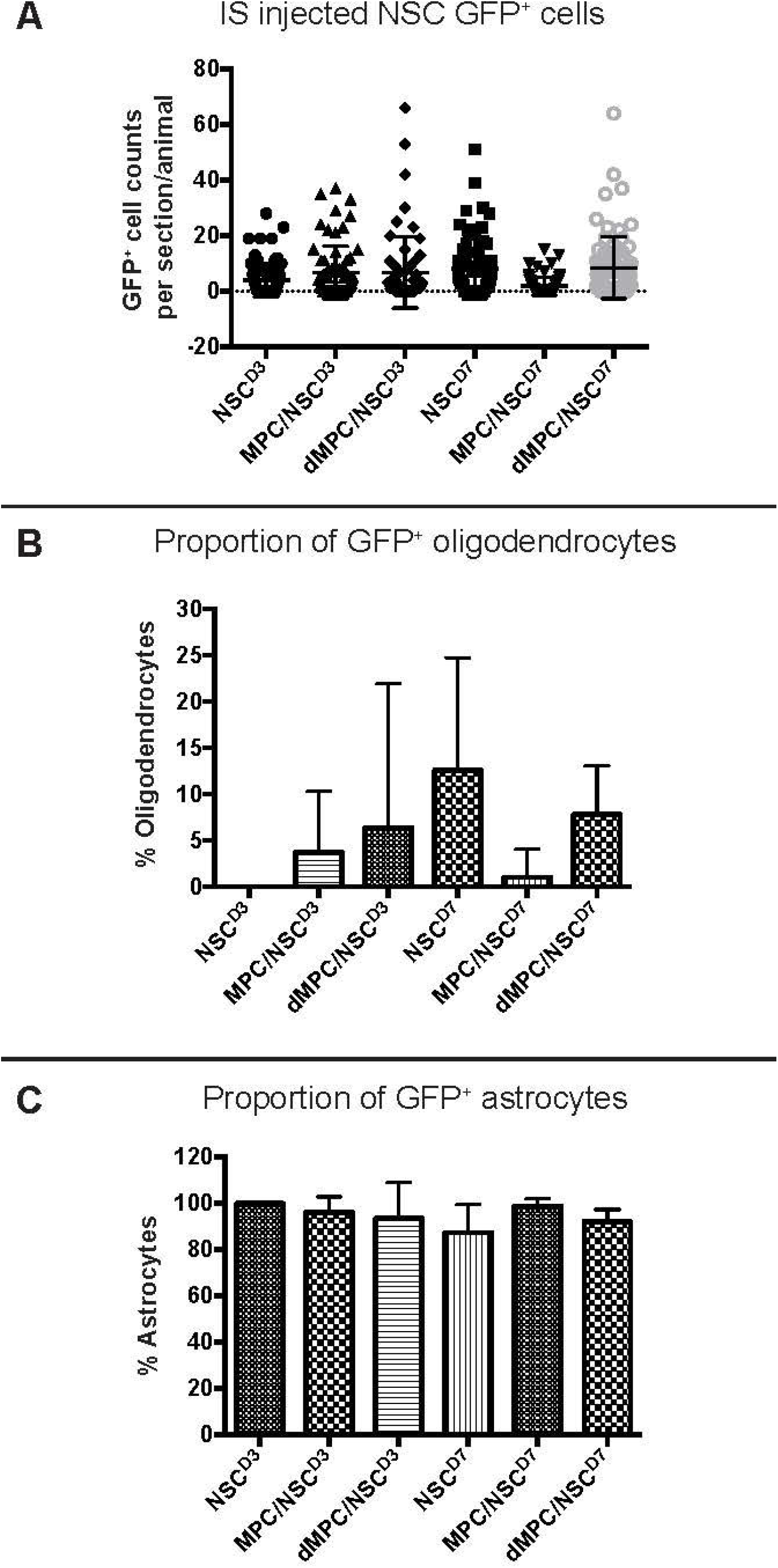
GFP+ NSC cell counts in the spinal cord 6 weeks after injury. (A) The number of GFP+ NSCs in the processed spinal cord tissue was counted per section per mouse and displayed as a dot plot summary. (B) The number of GFP+ NSCs that differentiated into oligodendrocytes was counted and plotted as a percentage from the total cells. (C) The number of GFP+ NSCs that differentiated into astrocytes was counted and plotted as a percentage from the total cells. The percentage of oligodendrocytes and astrocytes are inversely proportional as no transplanted NSCs differentiated into neurons. There was no statistical significance seen in any of the groups.

When NSCs were injected at D3, approximately 5% of the total surviving NSCs differentiated into oligodendrocytes in the dMPC_D1/NSC_D3 group. For the MPC_D1/NSC_D3 group, approximately 3% of the total surviving NSCs differentiated into oligodendrocytes. However, only 25% (2 in 8) of the dMPC_D1/NSC_D3 mice had any oligodendrocytes. Of note, one mouse in the dMPC_D1/NSC_D3 had robust oligodendrocyte differentiation (Figure 7C). This was not seen in any other treatment group with or without intravenous injection. For the MPC_D1/NSC_D3 group, 44.4% (4 in 9) of the mice had NSCs differentiating into oligodendrocytes. In the NSCD3 group, 0% (0 in 9) of the mice had NSCs differentiating into oligodendrocytes.

**Figure 7.**
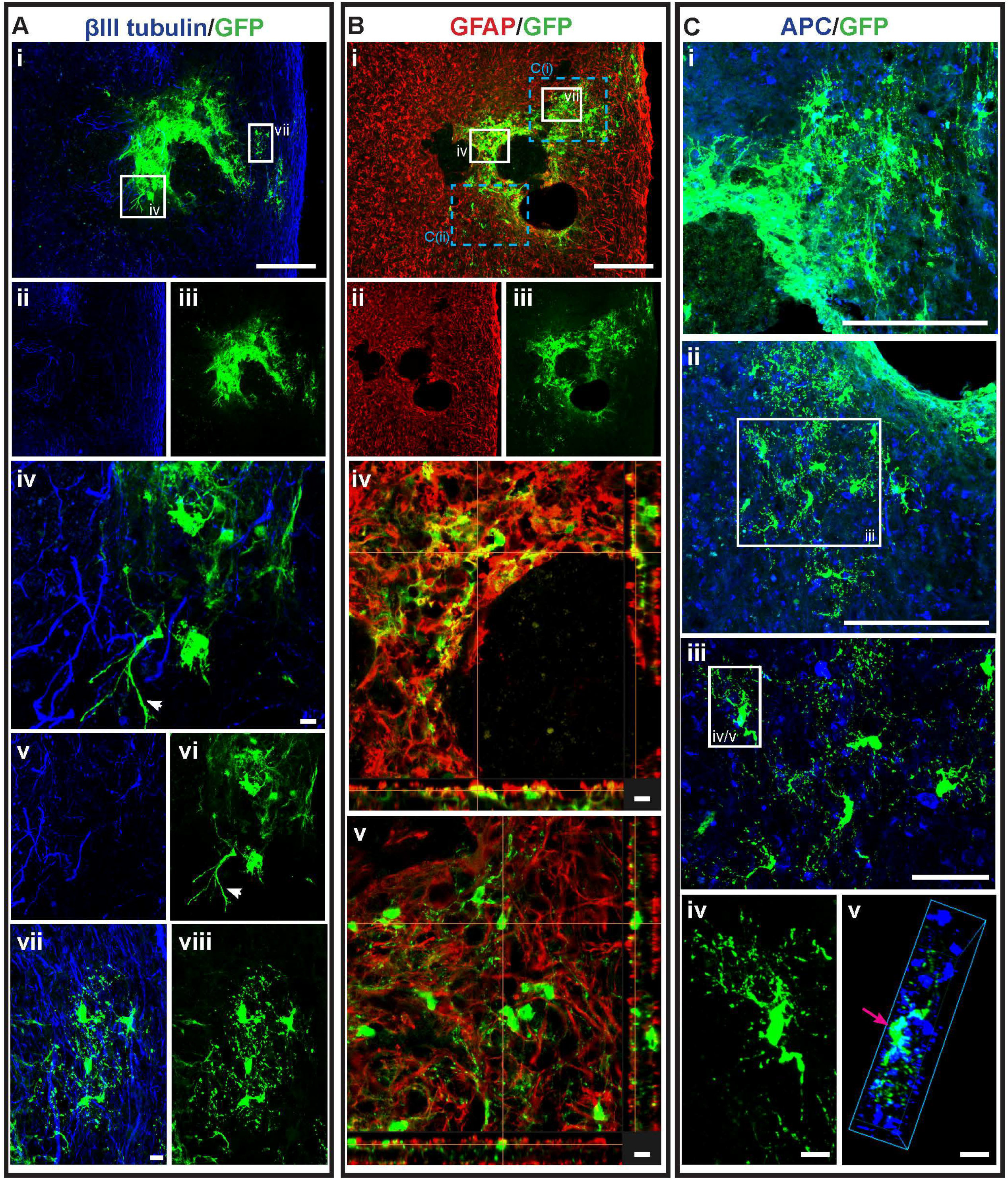
GFP+ NSCs in spinal cord sections of dMPC_D1/NSC_D3 mice. Gross over-view of surviving GFP+ NSCs transplanted at day 3 after intravenous injection of dead MPCs at D1 following cervical spinal cord injury showing (Ai, ii, iii) βIII-tubulin and GFP. High magnification images as outlined in the white boxes are shown in iv-viii as follows: (Aiv) GFP+ astrocytes with βIII-tubulin, (Av) βIII-tubulin+ axonal profiles, and (Avi) GFP+ astrocytes, (Avii) GFP+ oligodendrocytes with βIII-tubulin and (Aviii) GFP+ oligodendrocytes. The white arrows seen in Aiv and Avi shows an astrocyte projecting long processes. Astrocytic profile of the spinal cord tissue around the injury site is seen in (Bi, ii, iii) showing GFAP+ astrocytes and GFP+ differentiated NSCs. (Biv) and (Bv) shows high magnification image of the area outline in the white boxes in Bi. The orange lines intersecting the images shows the dual-label of GFAP and GFP on the cell that the line intersects in the vertical column on the right and horizontal column underneath. The blue dotted boxes in Bi corresponds to the figures in (Ci) and (Cii) showing oligodendrocyte profile of the spinal cord tissue around the injury site with APC+ oligodendrocytes and GFP+ differentiated NSCs. The high magnification image of the area in the white box in Cii is shown in (Ciii). (Civ) and shows the GFP+ labeled oligodendrocyte outlined by the white box in Ciii. (Cv) is the 3D view of the same cell as shown by the pink arrow. Scale bars for Ai, Bi, Ci, Cii = 200 μm, Ciii = 50 μm, Aiv, Avii, Biv, Bv, Civ, Cv = 10 μm.

For the NSC D7 treatment groups, mice receiving dMPC_D1/NSC_D7showed increased oligodendrocyte differentiation compared to mice receiving MPC_D1/NSC_D7with approximately 8% of the total surviving NSCs differentiating into oligodendrocytes versus <1% (Figure 6B). More notably, 100% (10 in 10) of the dMPC_D1/NSC_D7treated mice had intraspinal injected NSCs differentiate into oligodendrocytes whereas only 11.1% (1 in 9) MPC_D1/NSC_D7treated mice had intraspinal injected NSCs differentiate into oligodendrocytes. In the NSCD7 group, 75% (6 in 8) of the mice had transplanted NSCs that differentiated into oligodendrocytes. Differentiated GFP+ oligodendrocytes in the dMPC_D1/NSC_D7groups were generally found in clusters of at least 3-4 cells (Figure 7, 8) and moved in a rostral or caudal direction to the injection site rather than to the periphery as seen in the MPC_D1/NSC_D7or NSCD7 groups.

**Figure 8.**
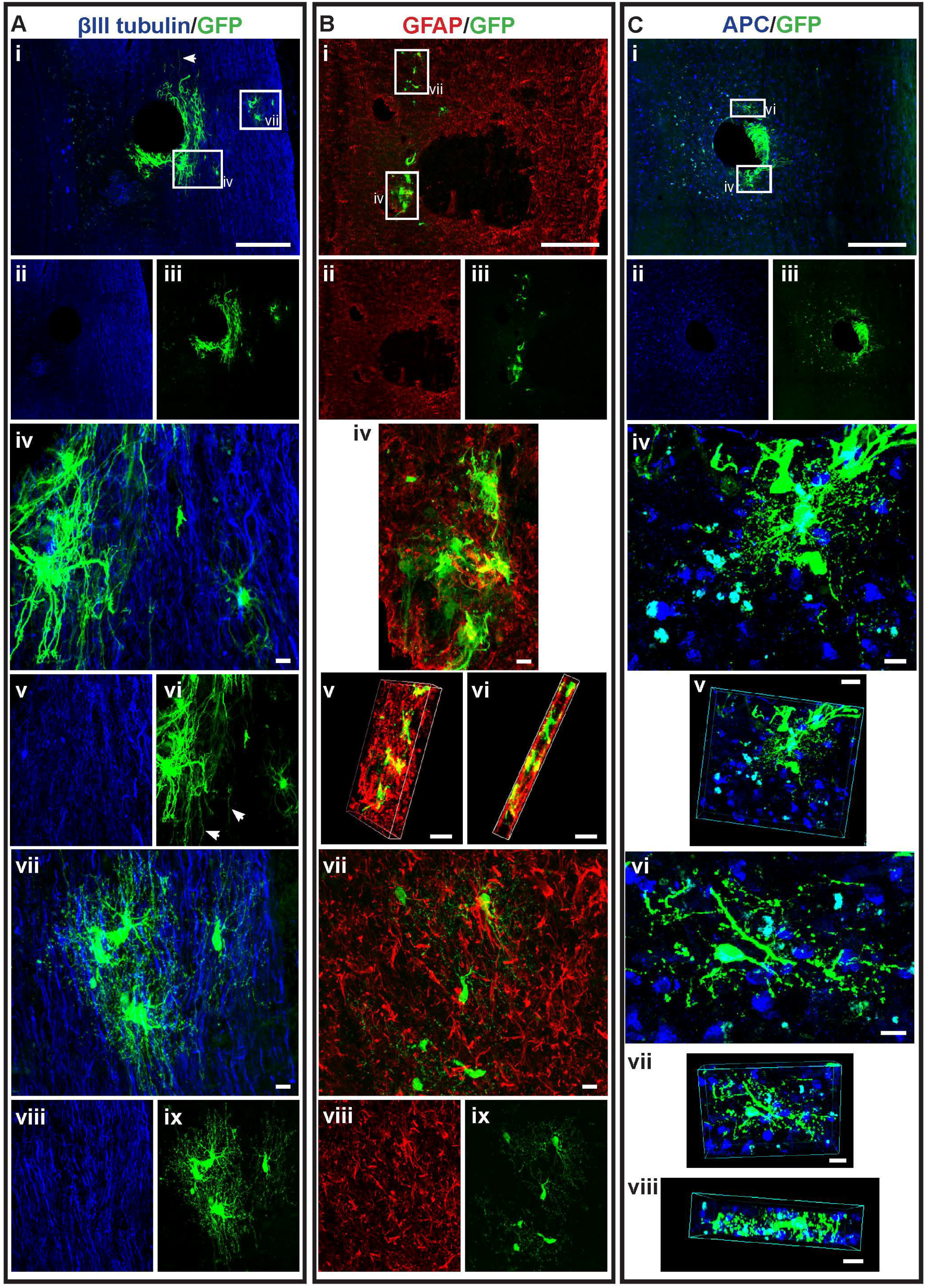
GFP+ NSCs in spinal cord sections of dMPC_D1/NSC_D7 mice. Gross over-view of surviving GFP+ NSCs transplanted at day 7 after intravenous injection of dead MPCs at D1 following cervical spinal cord injury showing (Ai, ii, iii) βIII-tubulin and GFP. High magnification images as outlined in the white boxes are shown in iv-viii as follows: (Aiv) GFP+ astrocytes with βIII-tubulin, (Av) βIII-tubulin+ axonal profiles, and (Avi) GFP+ astrocytes, (Avii) GFP+ oligodendrocytes with βIII-tubulin, (Aviii) βIII-tubulin+ axonal profiles and (Aix) GFP+ oligodendrocytes. The white arrows seen in Aiv show astrocytes projecting long processes. Astrocytic profile of the spinal cord tissue around the injury site is seen in (Bi, ii, iii) showing GFAP+ astrocytes and GFP+differentiated NSCs. (Biv) shows high magnification image of the area outline in the white box in Bi. (Bv) and (Bvi) shows the 3D view of the same image in Biv showing dual-label of cells with GFAP and GFP. (Bvii) shows the dual-labeled high magnification image of the area outline in the white box in Bi while the single channel image is seen in (Bviii) for GFAP+ axonal profile and (Bix) for GFP+ oligodendrocytes. Oligodendrocyte profile of the spinal cord tissue around the injury site is seen in (Ci, ii, iii) showing APC+ oligodendrocytes and GFP+ differentiated NSCs. (Civ) shows the dual-labeled high magnification image of the area outline in the white box in Ci while (Cv) shows the 3D view of the same area. (Cvi) shows the dual-labeled high magnification images of the area outline in the white box in Ci while (Cvii) and (Cviii) show the 3D view of the same area. Scale bars for Ai, Bi, Ci, Cii = 200 μm while the rest of the scale bars = 10 μm.

### Dual-transplantation study: Intravenous injection of dead MPCs result in transplanted NSCs differentiating into astrocytes with long processes

Another notable characteristic in the differentiated NSCs in mice receiving IV injection of dead MPCs was that a small number of GFP+ astrocytes that possessed elongated processes (Figure 7, 8 - white arrows, Figure 9). This was not seen in any of the sections in the mice receiving MPCs and regardless of whether NSCs were injected at D3 or D7 after IV injection of dMPCs.

**Figure 9.**
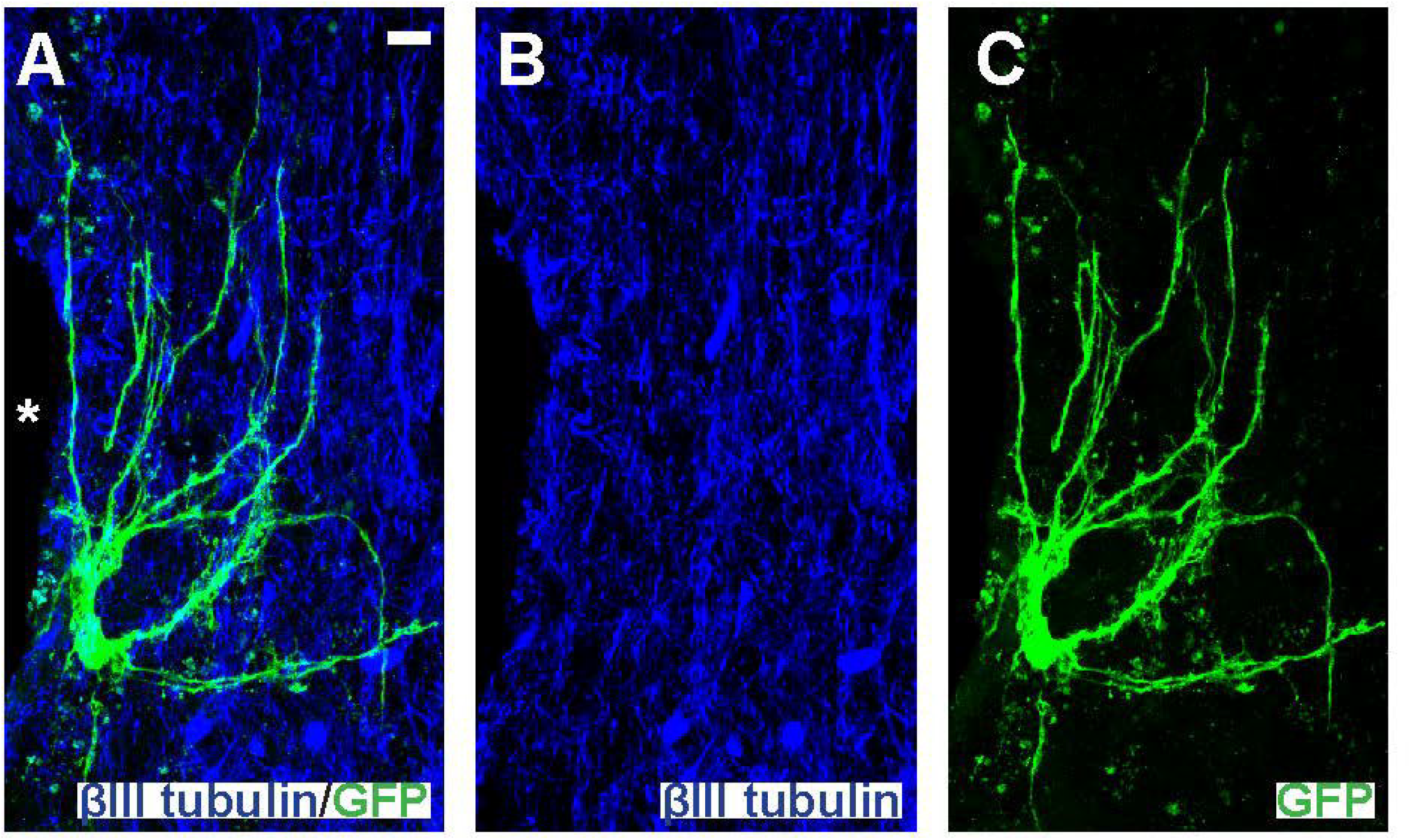
Intraspinal injected NSCs differentiated into astrocytes with long processes. High magnification image of an area around injection site (^*^) in a mouse receiving dMPC_D1/NSC_D7showing differentiated NSC into astrocyte projecting long processes with (A) dual-label of βIII-tubulin and GFP, (B) for single channel view of βIII-tubulin and (C) single channel view of GFP. Scale bar = 10 μm.

## 4. Discussion

The present study aimed to examine if heat treated (70°C) MPCs are a viable cell transplant control for SCI or do they elicit any therapeutic effects in the injured spinal cord. We compared the results from single IV injections of dMPCs versus IV injections MPCs or HBSS vehicle control, and in a dual-injection study when a single IV injection of dMPCs or MPCs was delivered at D1 post-injury followed by intraspinal injection of NSCs at D3 or D7 post-injury. In all studies, a cervical contusion injury model was used. Overall, our results demonstrated the following: (1) dMPCs obtained through prolonged heat treatment elicit positive results which effectively rejects using dMPCs as a cellular control for MPC injections, (2) dMPCs are not as effective as MPCs in promoting repair of the injured spinal cord, but had measurable positive results when compared to the vehicle control. (3) Functional and anatomical data from mice receiving dMPC injections was highly variable within each experimental group.

An issue with using dMPCs as a cellular control was the ability to obtain an injectable cell suspension that contained intact dMPCs in 300 μL of HBSS. Previous studies have shown that freeze thawing of grafts can produce intact grafts for transplantation [39, 40, 42], but this was not the case for cell suspensions. We had initially attempted to produce dead cells by rapidly freeze-thawing the MPCs cell suspension [40]. However, the resulting cell suspension was unusable as it resulted in the cells becoming lysed, membrane disintegration and DNA leakage, resulting in a sticky suspension. This then led to the use of heat for obtaining dMPCs.

While it was possible to obtain intact dMPCs through prolonged heating for IV injections, attempts to isolate the dMPCs from the supernatant in which cell death process occurred resulted in intact membranes fragmenting after centrifugation. Therefore, the HBSS media in which the cells were slowly killed over 12 hours was the same media used for injections. Due to the way dead cells were obtained, it is possible that transcription factors and surface markers related to cell death was secreted and maintained in the suspension. The inclusion of such factors could account for the positive results found when dMPCs was used compared to the vehicle control. The presence of different secreted factors resultant from the cell death process could also account for the variation in the results seen in the dual-injection study where some mice had more robust oligodendrocytes differentiation compared to others within the dMPCs group. This is further supported by studies that used conditioned medium from NSCs and bone-marrow derived mesenchymal cells administered in rats with SCI showing anatomical and functional improvements [48, 49]. These studies showed that the media itself contains factors to help promote repair after SCI. Therefore, dMPCs obtained through prolonged heating in this study is unsuitable as a cellular control for MPCs in cervical SCI.

MPCs have historically been shown to secrete immunomodulatory and trophic molecules [50-53]. In vivo, MPCs modulate immune responses through the regulation of the proliferation, recruitment, and the activation of innate and adaptive immune cells, such as T cells, B cells, dendritic cells, natural killer T cells, and macrophages [54, 55]. However, when MPCs experience stress, such as heat shock used here or oxidative stress, they can acquire a state of stressed induced premature senescence (SIPS) that changes their transcriptional pattern [56]. Such changes can even occur with a short burst of heat shock, Schlesinger and colleges [57] reported long term functional and transcriptional changes in MPCs following heat shock at 42°C for 1hr in vitro. SIPS MPCs can switch to senescent associated secretory profile (SASP), which includes proinflammatory cytokines such as IL1, IL6, and IL6 through a shift of transcriptional control of nuclear factor κB [58]. SASP can further induce DNA damage and senescence in neighboring cells [59]. Interestingly, heat shock to MPCs has been demonstrated to increase oxidative stress by increasing reactive oxygen species production or the downregulation of anti-oxidation functions [60-62]. In our context, prolonged heat shock at 72°C could induce MPCs into a senescent state before completely killing the cells and thus could include heat resistant secretory factors or reactive oxygen species in the supernatant. Possibly, proinflammatory molecules could act as a direction for the host immune response thus decreasing inflammation in the lesion core.

In our previous work, we suggested that the transplanted MPCs acted as a cellular target decoy by distracting the activated immune system away from the SCI site thereby resulting in a smaller lesion size and change in axonal, glia and vascularization profile compared to the vehicular control [29]. The distraction could occur through the physical presence of the MPC and/or through the factors secreted by the MPCs. By using dMPCs, the injected cell suspension could contain secretory factors produced by MPCs before complete cell death occurred. In addition, the physical presence of the dMPCs could have played a role in distracting the activated immune system after SCI thereby leading to the smaller lesion size and functional improvement seen. The response may not be as robust compared to the MPCs group as the secretory profile is limited in the dMPCs injection and the physical presence of dMPCs may not evoke as large a response from the innate immune system.

Previously, we have validated the presence of IV injected MPCs in the mice lungs following injection [29]. Recent studies have shown that MPCs modulate host immune responses following injury through extracellular vesicles (EVs) [63-66]. Outside of neurotrauma, EVs derived from MPCs has already been shown to reduce inflammation from bacterial infections [67, 68]. Further investigation to elucidate if this mechanism is indeed how MPCs modulate the distal lesion in the spinal cord. Similar with dMPCs, the release of EVs, namely apoptotic bodies (ApoBDs) or microvesicles (MV), in the process of the 12hour heat shock could facilitate apoptosis induced EVs release [69]. The released EVs could act as a distraction for the host immune response or alternatively release a similar secretome as live MPCs.

## 5. Conclusions

We conclude that dMPCs obtained through prolonged heat treatment elicits a positive outcome for MPCs transplantation studies in SCI compared to vehicle control, and thus would be inappropriate as a cellular control. It is possible that another method of preparation of dMPCs and/or an appropriately prepared dead cell of any other cell type can act as an appropriate cellular control for MPCs transplantation studies in SCI. If an appropriate cellular control can be found for transplantation studies, it will act as an invaluable tool to further elucidate the mechanisms in which transplanted cells can elicit repair on the injured spinal cord. Currently, it remains a challenge to identify a useable cellular control for cell transplantation studies in SCI.

## References

1. White SV, Ma YHE, Plant CD, Harvey AR, Plant GW. Combined Transplantation of Mesenchymal Progenitor and Neural Stem Cells to Repair Cervical Spinal Cord Injury. Cells. 2025;14(9):630. Published 2025 Apr 23. doi:10.3390/cells14090630

2. Akiyama Y, Honmou O, Kato T, Uede T, Hashi K, Kocsis JD. Transplantation of clonal neural precursor cells derived from adult human brain establishes functional peripheral myelin in the rat spinal cord. Exp Neurol. 2001;167(1):27–39. doi:10.1006/exnr.2000.7539

3. Ormond DR, Shannon C, Oppenheim J, et al. Stem cell therapy and curcumin synergistically enhance recovery from spinal cord injury. PLoS One. 2014;9(2):e88916. Published 2014 Feb 18. doi:10.1371/journal.pone.0088916

4. Poplawski GHD, Kawaguchi R, Van Niekerk E, et al. Injured adult neurons regress to an embryonic transcriptional growth state. Nature. 2020;581(7806):77–82. doi:10.1038/s41586-020-2200-5

5. Lu P, Freria CM, Graham L, et al. Rehabilitation combined with neural progenitor cell grafts enables functional recovery in chronic spinal cord injury. JCI Insight. 2022;7(16):e158000. Published 2022 Aug 22. doi:10.1172/jci.insight.158000

6. Doulames VM, Weimann JM, Plant GW. Human deep cortical neurons promote regeneration and recovery after cervical spinal cord injury. bioRxiv. 2021 doi: 10.1101/2021.08.11.455948.

7. Doulames VM, Marquardt LM, Hefferon ME, Baugh NJ, Suhar RA, Wang AT, Dubbin KR, Weimann JM, Palmer TD, Plant GW, Heilshorn SC. Custom-engineered hydrogels for delivery of human hiPSC-derived neurons into the injured cervical spinal cord. Biomaterials. 2024;305:122400. doi: 10.1016/j.biomaterials.2023.122400.

8. Olmsted ZT, Stigliano C, Marzullo B, Cibelli J, Horner PJ, Paluh JL. Fully characterized mature human iPS- and NMP-drived motor neurons thrive without neuroprotection in the spinal contusion cavity. Front Cell Neurosci. 2022;15:725195. doi: 10.3389/fncel.2021.725195.

9. Zholudeva LV, Fortino T, Agrawal A, Vila OF, Williams M, McDevitt T, Lane MA, Srivastava D. Human spinal inter-neurons repair the injured spinal cord through synaptic integration. bioRxiv [preprint] 2024 doi: 10.1101/2024.01.11.575264.

10. Kanno H, Pearse DD, Ozawa H, Itoi E, Bunge MB. Schwann cell transplantation for spinal cord injury repair: its significant therapeutic potential and prospectus. Rev Neurosci. 2015;26(2):121–128. doi:10.1515/revneuro-2014-0068

11. Marquardt LM, Doulames VM, Wang AT, et al. Designer, injectable gels to prevent transplanted Schwann cell loss during spinal cord injury therapy. Sci Adv. 2020;6(14):eaaz1039. Published 2020 Apr 1. doi:10.1126/sciadv.aaz1039

12. Coutts DJ, Humphries CE, Zhao C, Plant GW, Franklin RJ. Embryonic-derived olfactory ensheathing cells remyelinate focal areas of spinal cord demyelination more efficiently than neonatal or adult-derived cells. Cell Transplant. 2013;22(7):1249–1261. doi:10.3727/096368912X656153

13. Awidi A, Al Shudifat A, El Adwan N, et al. Safety and potential efficacy of expanded mesenchymal stromal cells of bone marrow and umbilical cord origins in patients with chronic spinal cord injuries: a phase I/II study. Cytotherapy. 2024;26(8):825–831. doi:10.1016/j.jcyt.2024.03.480

14. Vawda R, Badner A, Hong J, et al. Early Intravenous Infusion of Mesenchymal Stromal Cells Exerts a Tissue Source Age-Dependent Beneficial Effect on Neurovascular Integrity and Neurobehavioral Recovery After Traumatic Cervical Spinal Cord Injury. Stem Cells Transl Med. 2019;8(7):639–649. doi:10.1002/sctm.18-0192

15. Yao S, Pang M, Wang Y, et al. Mesenchymal stem cell attenuates spinal cord injury by inhibiting mitochondrial quality control-associated neuronal ferroptosis. Redox Biol. 2023;67:102871. doi:10.1016/j.redox.2023.102871

16. Barbour, H.R., Plant, C.D., Harvey, A.R., Plant, G.W., 2013. Tissue sparing, behavioral recovery, supraspinal axonal sparing/regeneration following sub-acute glial transplantation in a model of spinal cord contusion. BMC neuroscience 14, 106.

17. Li, B.C., Xu, C., Zhang, J.Y., Li, Y., Duan, Z.X., 2012. Differing Schwann cells and olfactory ensheathing cells behaviors, from interacting with astrocyte, produce similar improvements in contused rat spinal cord’s motor function. Journal of molecular neuroscience : MN 48, 35–44.

18. Plant, G.W., Christensen, C.L., Oudega, M., Bunge, M.B., 2003. Delayed transplantation of olfactory ensheathing glia promotes sparing/regeneration of supraspinal axons in the contused adult rat spinal cord. Journal of neurotrauma 20, 1–16.

19. Takami, T., Oudega, M., Bates, M.L., Wood, P.M., Kleitman, N., Bunge, M.B., 2002. Schwann cell but not olfactory ensheathing glia transplants improve hindlimb locomotor performance in the moderately contused adult rat thoracic spinal cord. The Journal of neuroscience : the official journal of the Society for Neuroscience 22, 6670–6681.

20. Takami, T., Oudega, M., Bates, M.L., Wood, P.M., Kleitman, N., Bunge, M.B., 2002. Schwann cell but not olfactory ensheathing glia transplants improve hindlimb locomotor performance in the moderately contused adult rat thoracic spinal cord. The Journal of neuroscience : the official journal of the Society for Neuroscience 22, 6670–6681.

21. Xu, X.M., Chen, A., Guenard, V., Kleitman, N., Bunge, M.B., 1997. Bridging Schwann cell transplants promote axonal regeneration from both the rostral and caudal stumps of transected adult rat spinal cord. Journal of neurocytology 26, 1–16.

22. Barraud, P., Seferiadis, A.A., Tyson, L.D., Zwart, M.F., Szabo-Rogers, H.L., Ruhrberg, C., Liu, K.J., Baker, C.V., 2010. Neural crest origin of olfactory ensheathing glia. Proceedings of the National Academy of Sciences of the United States of America 107, 21040–21045.

23. Hodgetts, S.I., Simmons, P.J., Plant, G.W., 2013. Human mesenchymal precursor cells (Stro-1(+)) from spinal cord injury patients improve functional recovery and tissue sparing in an acute spinal cord injury rat model. Cell transplantation 22, 393–412.

24. Park, J.H., Min, J., Baek, S.R., Kim, S.W., Kwon, I.K., Jeon, S.R., 2013. Enhanced neuroregenerative effects by scaffold for the treatment of a rat spinal cord injury with Wnt3a-secreting fibroblasts. Acta neurochirurgica 155, 809–816.

25. Tetzlaff, W., Okon, E.B., Karimi-Abdolrezaee, S., Hill, C.E., Sparling, J.S., Plemel, J.R., Plunet, W.T., Tsai, E.C., Baptiste, D., Smithson, L.J., Kawaja, M.D., Fehlings, M.G., Kwon, B.K., 2011. A systematic review of cellular transplantation therapies for spinal cord injury. Journal of neurotrauma 28, 1611–1682.

26. Jin, Y., Neuhuber, B., Singh, A., Bouyer, J., Lepore, A., Bonner, J., Himes, T., Campanelli, J.T., Fischer, I., 2011. Transplantation of human glial restricted progenitors and derived astrocytes into a contusion model of spinal cord injury. Journal of neurotrauma 28, 579–594.

27. Yu, T.B., Cheng, Y.S., Zhao, P., Kou, D.W., Sun, K., Chen, B.H., Wang, A.M., 2009. Immune therapy with cultured microglia grafting into the injured spinal cord promoting the recovery of rat’s hind limb motor function. Chinese journal of traumatology = Zhonghua chuang shang za zhi /Chinese Medical Association 12, 291–295.

28. Ma, S.F., Chen, Y.J., Zhang, J.X., Shen, L., Wang, R., Zhou, J.S., Hu, J.G., Lu, H.Z., 2015. Adoptive transfer of M2 macrophages promotes locomotor recovery in adult rats after spinal cord injury. Brain, behavior, and immunity 45, 157–170.

29. White, S.V., Czisch, C.E., Han, M.H., Plant, C.D., Harvey, A.R., Plant, G.W., 2016. Intravenous Transplantation Of Mesenchymal Progenitors Distribute Solely To The Lungs And Improve Outcomes in Cervical Spinal Cord Injury. Stem cells.

30. Haase, S.C., Rovak, J.M., Dennis, R.G., Kuzon, W.M., Jr., Cederna, P.S., 2003. Recovery of muscle contractile function following nerve gap repair with chemically acellularized peripheral nerve grafts. Journal of reconstructive microsurgery 19, 241–248.

31. Hudson, T.W., Liu, S.Y., Schmidt, C.E., 2004. Engineering an improved acellular nerve graft via optimized chemical processing. Tissue engineering 10, 1346–1358.

32. Mligiliche, N., Kitada, M., Ide, C., 2001. Grafting of detergent-denatured skeletal muscles provides effective conduits for extension of regenerating axons in the rat sciatic nerve. Archives of histology and cytology 64, 29–36.

33. Evans, P.J., Mackinnon, S.E., Levi, A.D., Wade, J.A., Hunter, D.A., Nakao, Y., Midha, R., 1998. Cold preserved nerve allografts: changes in basement membrane, viability, immunogenicity, and regeneration. Muscle & nerve 21, 1507–1522.

34. Ide, C., 1983. Nerve regeneration and Schwann cell basal lamina: observations of the long-term regeneration. Archivum histologicum Japonicum = Nihon soshikigaku kiroku 46, 243–257.

35. Ide, C., Tohyama, K., Yokota, R., Nitatori, T., Onodera, S., 1983. Schwann cell basal lamina and nerve regeneration. Brain research 288, 61–75.

36. Osawa, T., Ide, C., Tohyama, K., 1987. Nerve regeneration through cryo-treated xenogeneic nerve grafts. Archivum histologicum Japonicum = Nihon soshikigaku kiroku 50, 193–208.

37. Mackinnon, S.E., Hudson, A.R., Falk, R.E., Kline, D., Hunter, D., 1984a. Peripheral nerve allograft: an assessment of regeneration across pretreated nerve allografts. Neurosurgery 15, 690–693.

38. Mackinnon, S.E., Hudson, A.R., Falk, R.E., Kline, D., Hunter, D., 1984b. Peripheral nerve allograft: an immunological assessment of pretreatment methods. Neurosurgery 14, 167–171.

39. Enver, M.K., Hall, S.M., 1994. Are Schwann cells essential for axonal regeneration into muscle autografts? Neuropathology and applied neurobiology 20, 587–598.

40. Cui, Q., Pollett, M.A., Symons, N.A., Plant, G.W., Harvey, A.R., 2003. A new approach to CNS repair using chimeric peripheral nerve grafts. Journal of neurotrauma 20, 17–31.

41. Bamber, N.I., Li, H., Aebischer, P., Xu, X.M., 1999. Fetal spinal cord tissue in mini-guidance channels promotes longitudinal axonal growth after grafting into hemisected adult rat spinal cords. Neural plasticity 6, 103–121.

42. Hall, S.M., Enver, K., 1994. Axonal regeneration through heat pretreated muscle autografts. An immunohistochemical and electron microscopic study. Journal of hand surgery 19, 444–451.

43. Cao, Y.A., Wagers, A.J., Beilhack, A., Dusich, J., Bachmann, M.H., Negrin, R.S., Weissman, I.L., Contag, C.H., 2004. Shifting foci of hematopoiesis during reconstitution from single stem cells. Proceedings of the National Academy of Sciences of the United States of America 101, 221–226.

44. van der Bogt, K.E., Hellingman, A.A., Lijkwan, M.A., Bos, E.J., de Vries, M.R., van Rappard, J.R., Fischbein, M.P., Quax, P.H., Robbins, R.C., Hamming, J.F., Wu, J.C., 2012. Molecular imaging of bone marrow mononuclear cell survival and homing in murine peripheral artery disease. JACC. Cardiovascular imaging 5, 46–55.

45. Short, B.J., Brouard, N., Simmons, P.J., 2009. Prospective isolation of mesenchymal stem cells from mouse compact bone. Methods in molecular biology 482, 259–268.

46. Azari, H., Rahman, M., Sharififar, S., Reynolds, B.A., 2010. Isolation and expansion of the adult mouse neural stem cells using the neurosphere assay. Journal of visualized experiments : JoVE.

47. Schallert, T., Fleming, S.M., Leasure, J.L., Tillerson, J.L., Bland, S.T., 2000. CNS plasticity and assessment of forelimb sensorimotor outcome in unilateral rat models of stroke, cortical ablation, parkinsonism and spinal cord injury. Neuropharmacology 39, 777–787.

48. Cantinieaux, D., Quertainmont, R., Blacher, S., Rossi, L., Wanet, T., Noel, A., Brook, G., Schoenen, J., Franzen, R., 2013. Conditioned medium from bone marrow-derived mesenchymal stem cells improves recovery after spinal cord injury in rats: an original strategy to avoid cell transplantation. PloS one 8, e69515.

49. Liang, P., Liu, J., Xiong, J., Liu, Q., Zhao, J., Liang, H., Zhao, L., Tang, H., 2014. Neural stem cell-conditioned medium protects neurons and promotes propriospinal neurons relay neural circuit reconnection after spinal cord injury. Cell transplantation 23 Suppl 1, S45–56.

50. Carrade DD, Lame MW, Kent MS, Clark KC, Walker NJ, Borjesson DL. Comparative Analysis of the Immunomodulatory Properties of Equine Adult-Derived Mesenchymal Stem Cells(). Cell Med. 2012;4(1):1–11. doi:10.3727/215517912X647217

51. Kuroda Y, Dezawa M. Mesenchymal stem cells and their subpopulation, pluripotent muse cells, in basic research and regenerative medicine. Anat Rec (Hoboken). 2014;297(1):98–110. doi:10.1002/ar.22798

52. Li YW, Zhang C, Sheng QJ, Bai H, Ding Y, Dou XG. Mesenchymal stem cells rescue acute hepatic failure by polarizing M2 macrophages. World J Gastroenterol. 2017;23(45):7978–7988. doi:10.3748/wjg.v23.i45.7978

53. Shi Y, Wang Y, Li Q, et al. Immunoregulatory mechanisms of mesenchymal stem and stromal cells in inflammatory diseases. Nat Rev Nephrol. 2018;14(8):493–507. doi:10.1038/s41581-018-0023-5

54. Aggarwal S, Pittenger MF. Human mesenchymal stem cells modulate allogeneic immune cell responses. Blood. 2005;105(4):1815–1822. doi:10.1182/blood-2004-04-1559

55. Squillaro T, Severino V, Alessio N, et al. De-regulated expression of the BRG1 chromatin remodeling factor in bone marrow mesenchymal stromal cells induces senescence associated with the silencing of NANOG and changes in the levels of chromatin proteins. Cell Cycle. 2015;14(8):1315–1326. doi:10.4161/15384101.2014.995053

56. de Magalhães JP, Passos JF. Stress, cell senescence and organismal ageing. Mech Ageing Dev. 2018;170:2–9. doi:10.1016/j.mad.2017.07.001

57. Ribarski-Chorev I, Schudy G, Strauss C, Schlesinger S. Short heat shock has a long-term effect on mesenchymal stem cells’ transcriptome. iScience. 2023;26(8):107305. Published 2023 Jul 10. doi:10.1016/j.isci.2023.107305

58. Coppé JP, Patil CK, Rodier F, et al. Senescence-associated secretory phenotypes reveal cell-nonautonomous functions of oncogenic RAS and the p53 tumor suppressor. PLoS Biol. 2008;6(12):2853–2868. doi:10.1371/journal.pbio.0060301

59. Gorgoulis V, Adams PD, Alimonti A, et al. Cellular Senescence: Defining a Path Forward. Cell. 2019;179(4):813–827. doi:10.1016/j.cell.2019.10.005

60. Bernabucci U, Ronchi B, Lacetera N, Nardone A. Markers of oxidative status in plasma and erythrocytes of transition dairy cows during hot season. J Dairy Sci. 2002;85(9):2173–2179. doi:10.3168/jds.S0022-0302(02)74296-3

61. Abdelnour SA, Abd El-Hack ME, Khafaga AF, Arif M, Taha AE, Noreldin AE. Stress biomarkers and proteomics alteration to thermal stress in ruminants: A review. J Therm Biol. 2019;79:120–134. doi:10.1016/j.jtherbio.2018.12.013

62. Shimoni C, Goldstein M, Ribarski-Chorev I, et al. Heat Shock Alters Mesenchymal Stem Cell Identity and Induces Premature Senescence. Front Cell Dev Biol. 2020;8:565970. Published 2020 Sep 22. doi:10.3389/fcell.2020.565970

63. Nakazaki M, Morita T, Lankford KL, Askenase PW, Kocsis JD. Small extracellular vesicles released by infused mesenchymal stromal cells target M2 macrophages and promote TGF-β upregulation, microvascular stabilization and functional recovery in a rodent model of severe spinal cord injury. J Extracell Vesicles. 2021;10(11):e12137.

64. Liu W, Tang P, Wang J, et al. Extracellular vesicles derived from melatonin-preconditioned mesenchymal stem cells containing USP29 repair traumatic spinal cord injury by stabilizing NRF2. J Pineal Res. 2021;71(4):e12769. doi:10.1111/jpi.12769

65. Sun Y, Liu Q, Qin Y, et al. Exosomes derived from CD271+CD56+ bone marrow mesenchymal stem cell subpopoulation identified by single-cell RNA sequencing promote axon regeneration after spinal cord injury. Theranostics. 2024;14(2):510–527. Published 2024 Jan 1. doi:10.7150/thno.8900

66. Liu WZ, Ma ZJ, Li JR, Kang XW. Mesenchymal stem cell-derived exosomes: therapeutic opportunities and challenges for spinal cord injury. Stem Cell Res Ther. 2021;12(1):102. Published 2021 Feb 3. doi:10.1186/s13287-021-02153-8

67. Sarkar S, Barnaby R, Faber Z, et al. Extracellular Vesicles Derived from Mesenchymal Stromal Cells Reduce Pseudomonas aeruginosa Lung Infection and Inflammation in Mice. Preprint. bioRxiv. 2025;2025.03.30.646208. Published 2025 Mar 31. doi:10.1101/2025.03.30.646208

68. Sarkar S, Barnaby R, Nymon A, et al. Let-7b-5p loaded Mesenchymal Stromal Cell Extracellular Vesicles reduce Pseudomonas-biofilm formation and inflammation in CF Bronchial Epithelial Cells. Preprint. bioRxiv. 2025;2025.05.28.656674. Published 2025 May 28. doi:10.1101/2025.05.28.656674

69. Kakarla R, Hur J, Kim YJ, Kim J, Chwae YJ. Apoptotic cell-derived exosomes: messages from dying cells. Exp Mol Med. 2020;52(1):1–6. doi:10.1038/s12276-019-0362-8

